# Bladder-draining lymph nodes support germinal centre B cell responses during urinary tract infection in mice

**DOI:** 10.1101/2022.11.10.516078

**Authors:** Sophia Hawas, Dimitrios Vagenas, Ashraful Haque, Makrina Totsika

## Abstract

Bacterial urinary tract infections (UTIs) are both common and exhibit high recurrence rates in women. UTI healthcare costs are increasing due to the rise of multi-drug resistant (MDR) bacteria, necessitating alternative approaches for infection control. Here, we investigated whether host adaptive immune responses can influence infection outcomes. We employed a mouse model in which wild-type C57BL/6J mice were transurethrally inoculated with an MDR UTI strain of uropathogenic *Escherichia coli* (UPEC). Firstly, we noted that *rag1*^*-/-*^ C57BL/6J mice harboured larger bacterial burdens than wild-type counterparts, consistent with a role for T and/or B cells in optimal control of UTI. Consistent with this, UTI triggered in the bladders of wild-type mice early increases of myeloid cells, including CD11c^hi^ conventional dendritic cells, suggesting possible involvement of these professional antigen-presenting cells. Importantly, germinal centre (GC) B cell responses developed by 4 weeks post-infection in bladder-draining lymph nodes of wild-type mice, and although modest in magnitude and transient in nature, could not be boosted with a second UTI. Thus, our data reveal for the first time in a mouse model, that Gram-negative bacterial UTI induces local B cell immune responses in bladder-draining lymph nodes, which could potentially serve to control infection.

## Introduction

Urinary tract infections (UTIs) are one of the most common bacterial infections globally and contribute around US$6 billion to the annual global healthcare burden (1, 2). UTIs have a high rate of recurrence, over 30% of women with a primary infection will have a subsequent recurrent episode (1). Most UTIs are caused by uropathogenic *Escherichia coli* (UPEC) which are becoming increasingly multi-drug resistant bacteria (3). Drug resistance is a major problem in the treatment of UTIs, and a higher proportion of infections are now becoming recurrent, leading to complicated UTIs (4).

Currently, there is some evidence of adaptive immunity during UTI and the role it plays in affecting infection outcomes. Antigen-presenting cells such as dendritic cells are present during the inflammatory innate response, recruited by cytokines released by neutrophils and macrophages (5-7). Given their presence during innate immunity, it has been theorised that dendritic cells are responsible for ushering in the adaptive immune response and recruiting T and B cells to the bladder via antigen presentation and trafficking to local lymph nodes (5, 6, 8, 9). Previous studies in mouse knockout models of key immune cells (dendritic, T and B cells) have shown that adaptive immunity is important for multiple UTI outcomes, such as susceptibility to colonisation, bacterial clearance, and protection from subsequent infection (6, 10, 11). In a mouse model investigating the role of macrophages in UTI, Mora-Bau et al. found evidence of an adaptive immune response through recruited T and B cells to the bladder, which however did not offer sterilising immunity to infected mice (6). A key cytokine in innate UTI immunity is interleukin (IL)-17, the majority of which is generated by bladder-resident γδ T cells (10). Deficiency in this cytokine predisposes mice to bacterial persistence in the kidneys (10). In a γδ T cell-deficient mouse strain, mice had significantly higher bladder bacterial burdens compared to immunocompetent mice (11). A major component of adaptive immunity is the humoral response, which is responsible for antibody production and immunological memory. The cellular driver of the humoral response are B cells, which have been implicated to be important for sterilising immunity in UTIs (11, 12).

B cells are responsible for a wide variety of effector functions in humoral immunity, and naïve B cells require stimulation from an antigen, delivered to them by antigen-presenting cells (13). Naïve B cells in lymphatic tissue undergo maturation when exposed to an antigen, in a site within the lymph node known as the germinal centre. Here, they mature into activated germinal centre (GC) B cells which are the progenitor cell line for B cell subsets such as plasma and memory B cells (13), whose primary roles are to regulate antibody production. In UTI vaccination studies, experimental vaccines have been shown to elicit protective effects and stimulate antibody production in animal model infections (14-16). Antibody production has major implications for UTI clearance, and strong antibody responses are correlated with bacterial clearance (12). In a small cohort of human patients, few people with lower UTI had detectable antibody-secreting cells in their urine during the first few weeks of infection (14). In a monkey model of cystitis, anti-*E. coli* antibodies in urine peaked by 5 weeks post inoculation (wpi) (16), indicating that while the B cell response in UTI may not be as robust as in other infections, the contribution of these cells nonetheless is important for infection outcomes. Additionally, the presence of antibodies in serum and urine during acute UTI in both mouse and monkey models is associated with bacterial clearance and reduced bacterial numbers in urine (15, 16). These antibody responses arise from GC B cells, making them an important cellular target for UTI responses. To date, the contribution and dynamics of these GC B cells have not been explored in the mouse UTI model.

Here, we report the induction of B cell responses by observing GC B cells during acute UTI in mice, bridging studies of innate immunity and UTI vaccination. We first confirmed that T and B cells are important for infection outcomes and for controlling bacterial burden. We observed that GC B cells are generated in bladder draining lymph nodes during acute UTI, but are transient, and their population does not increase by secondary infection administered a week following the initial infection. Together, our study shows for the first time that humoral immunity is induced locally in bladder-draining lymph nodes in UTI and plays a role in controlling bacterial numbers, which has implications for UTI vaccine and drug development.

## Results

### Adaptive immunity contributes to acute UTI control

To investigate the contribution of adaptive immunity (T and B cells) to UPEC control during acute UTI, we compared bladder and urine bacterial loads in wild-type (C57BL/6) and *rag1*^*-/-*^ mice experimentally inoculated with the reference MDR ST131 UPEC strain EC958 (3) in the bladder. Colony forming units (CFU) in urine were collected over the course of the infection and analysed using a series of statistical models (including mixed, additive and zero inflated models) for parsimony, with the best fit being a Zero Inflated Negative Binomial Mixed Model (ZINBMM). At 4 wpi, *rag1*^*-/-*^ mice had 1-log higher median bacterial loads in the bladder compared to wild-type mice (*p* = 0.0017, unpaired non-parametric Mann-Whitney test) (Figure 1A). Urinalysis using ZINBMM also confirmed overall higher susceptibility for *rag1*^*-/-*^ mice, where modelling of urine data collected over time from three independent experiments, predicted a higher susceptibility to initial colonisation for *rag1*^*-/-*^ mice. However, once colonised, either mouse strain is predicted to have the same urinary bacterial burden over time. Combined, this results to an overall higher bacterial burden predicted for the *rag1*^*-/-*^ mouse group compared to WT mice (Figure 1B). Taken together, *rag1*^*-/-*^ mice are more susceptible to acute UTI, demonstrating a role for adaptive immunity in infection control.

**Figure 1.**
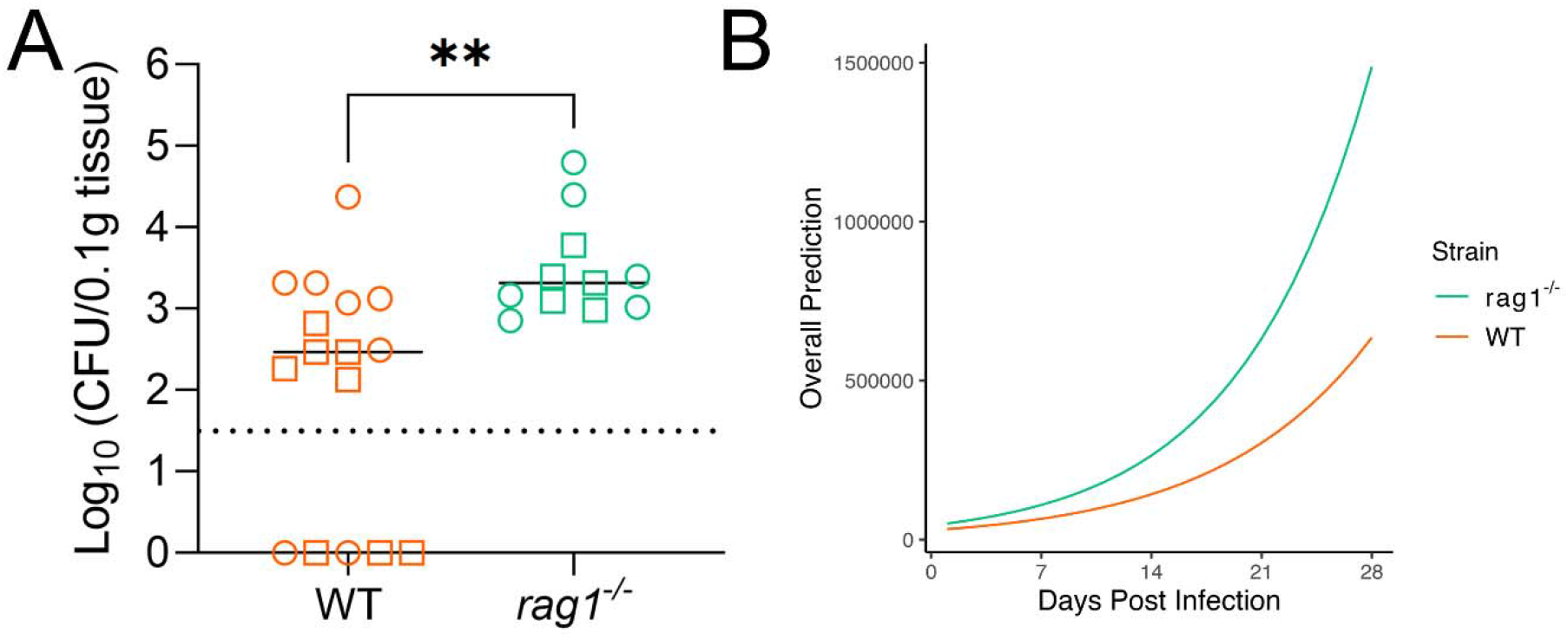
Impaired clearance of MDR UPEC from the *rag1^-/-^* mouse bladder. A) Scatter plot of C57BL/6 WT (*n* = 16) and *rag1*^*-/-*^ (*n* = 11) mouse bladder UPEC colonisation data (CFU/0.1g bladder tissue) at 4 weeks post inoculation (wpi) from two independent experiments. Lines represent group medians, dotted line represents limit of detection (LOD) and **\*\*** *p* < 0.01 (Mann-Whitney test). B) Longitudinal urinalysis showing the combined output of predicted colonisation chance and bacterial load in urine (CFU/mL urine over 28 days post infection) during UTI in WT (*n* = 26) and *rag1*^*-/-*^ mice (*n* = 11). Lines represent the overall predictions of a Zero Inflated Negative Binomial Mixed Model on urine data from three independent experiments.

### Dendritic and other innate immune cells are recruited to the bladder during acute infection

A robust innate immune response has been previously demonstrated in several UTI mouse studies (5, 7, 10, 15, 17-23). We confirmed this also occurs with our MDR ST131 UPEC strain, where after 24 hours of inoculation with EC958 we observed increased infiltration of monocytes (p=0.026), neutrophils (0.002), and dendritic cells (0.026) into the bladder of C57BL/6 WT mice in the UTI group compared to naïve controls (Figure 2B). Bladder and kidney bacterial loads at this timepoint were also assessed and found to be comparable to previous studies using the same mouse-UPEC strain combination (3) (data not shown).

**Figure 2.**
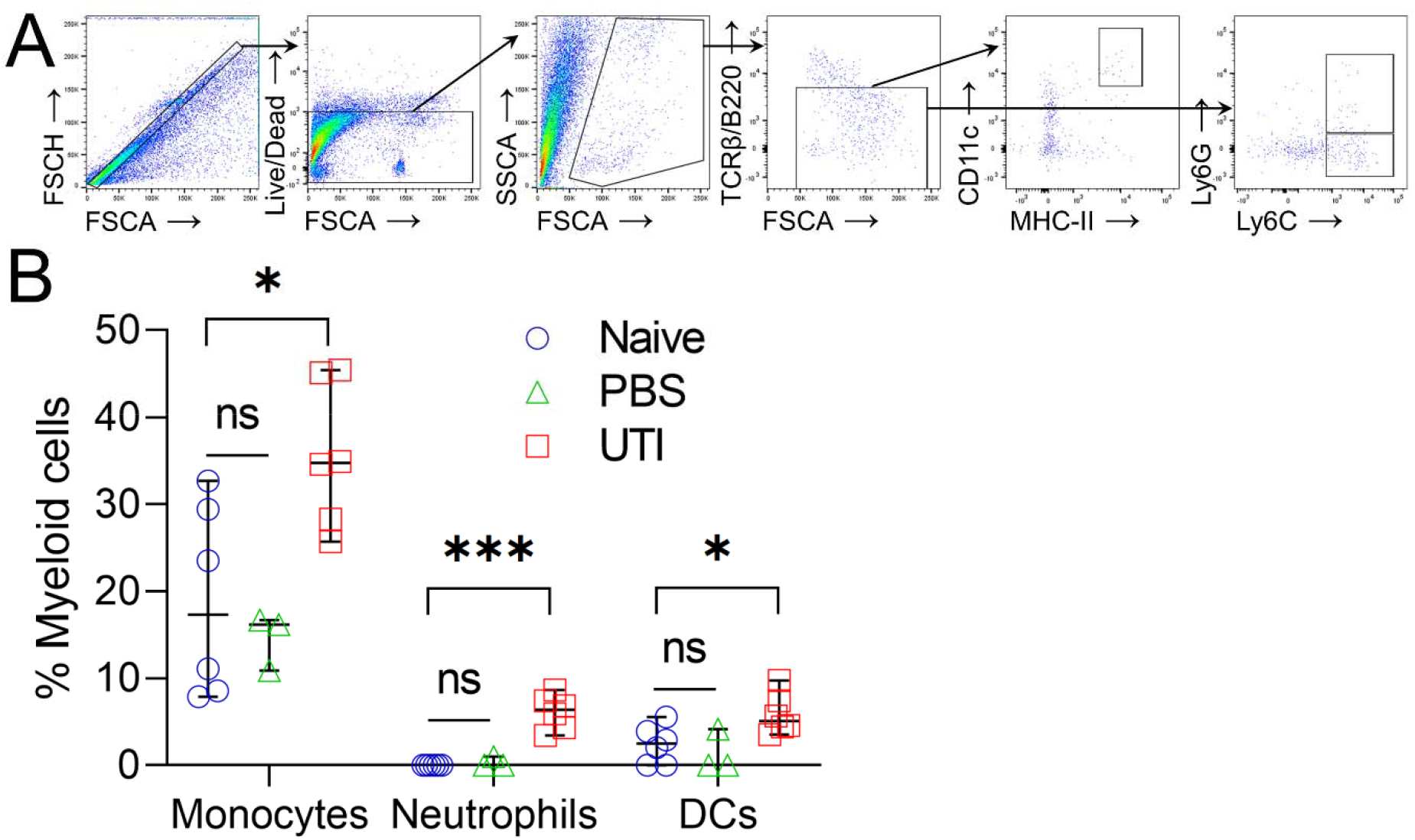
APCs (monocytes, neutrophils, and dendritic cells) are recruited to the C57BL/6 mouse bladder 1 day post inoculation with MDR UPEC. A) FACS gating strategy for myeloid cells (monocytes, neutrophils, and dendritic cells (DCs), B) Myeloid populations present in bladders of mice that were naive, transurethrally catheterised and inoculated with UPEC, or mock-catheterised with saline (PBS). Group differences detected by Kruskal-Wallis test (B), bars represent median ± 95% CI, **ns** not significant, **\*** *p* <0.05, **\*\*\*** *p* <0.001.

### B cell activation occurs locally and transiently during acute UTI

Despite the role of adaptive immunity in UTI, B cell subsets have not been previously reported in the UTI mouse model. Given the extensive investigation into the innate immune response in this model, specifically, the recruitment of dendritic cells to the bladder, we hypothesised that they would be responsible for triggering B cell responses in bladder-draining lymph nodes in UTI. We assessed bladder-draining lymph nodes of both naïve and inoculated mice at 4 weeks specifically to observe expected B cell responses at their peak (24). At 4 weeks post UPEC inoculation, we observed an increase in GC B cells in bladder-draining lymph nodes of C57BL/6 UTI mice (Figure 3C). While the proportion of GC B cells was relatively low (median 0.78%), this cell population was statistically larger than in the naïve mouse group (median 0.15%) (*p* <0.0001), providing supporting evidence for the induction of a local humoral response in UTI. However, we observed no differences in GC B cells in distal mesenteric lymph nodes (Figure 3D) or increases in plasmablast populations (IgD^lo^ CD138^+^) between groups (data not shown). This local response was also transient (Figure 3E), despite detectable CFU remaining present in the bladders of infected mice at 7 wpi (Figure 3F).

**Figure 3.**
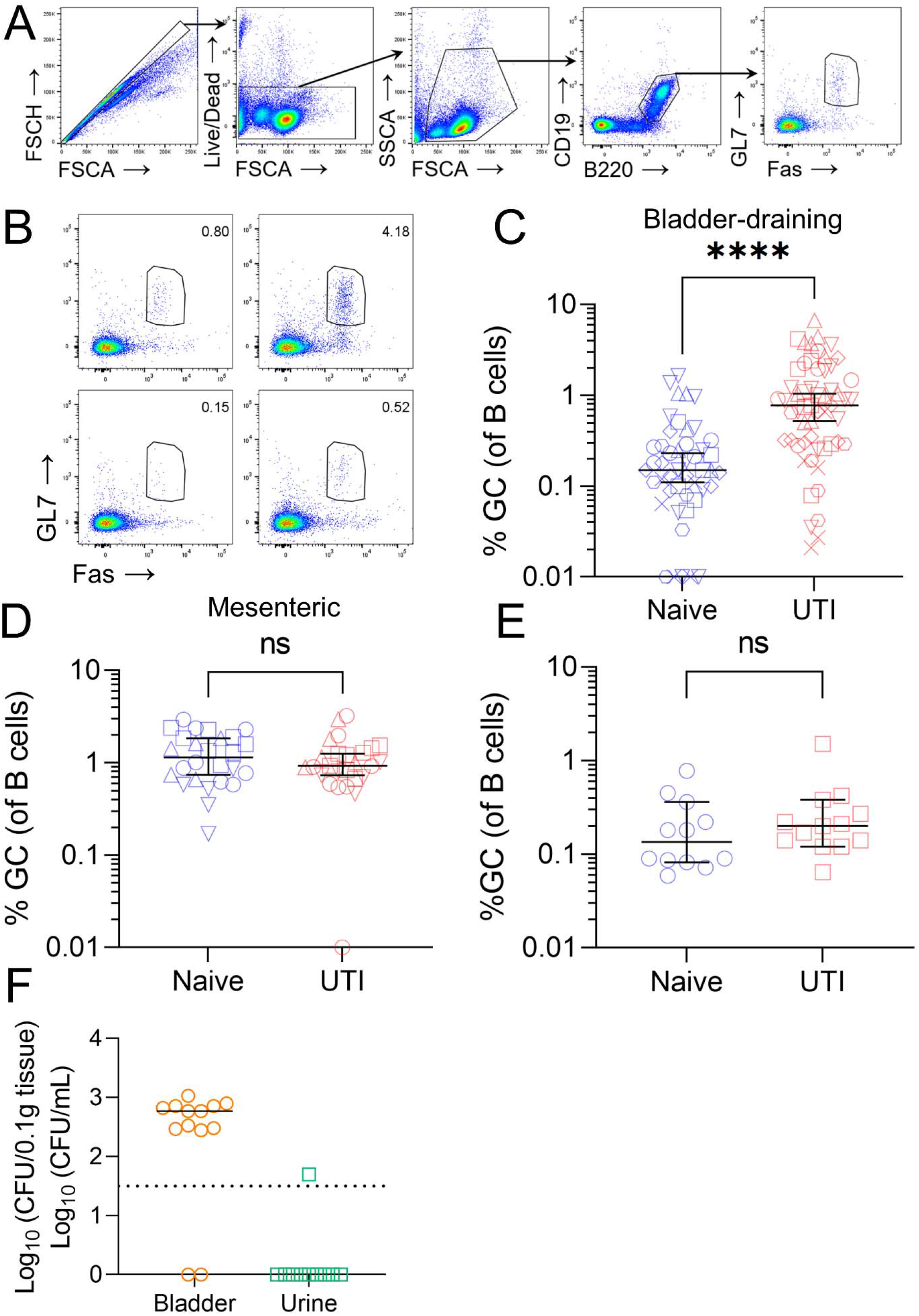
B cell activation occurs in bladder-draining lymph nodes at 4 weeks following acute UTI in C57BL/6 mice. A) FACS gating strategy for GC B cells and B) representative plots of the GC B cell gate in bladder-draining lymph nodes from two mice per group (UTI top, naïve bottom), % GC B cells (of total B cells detected) in C) bladder-draining lymph nodes at 4 wpi (UTI: *n* = 57, naïve: *n* = 47), D) mesenteric lymph nodes at 4 wpi (UTI: *n* = 26, naïve: *n* = 26), E) bladder-draining lymph nodes at 7 wpi (UTI: *n* = 13, naïve: *n* = 12), F) bacterial load in UTI mouse bladders (CFU/0.1g tissue) and urine (CFU/mL) at 7 wpi (*n* = 13). Lines in panels B-E represent group medians ±95% CI, and dotted lines represents LOD. Group difference detected by Mann-Whitney test, **ns** not significant, **\*\*\*\*** *p* <0.0001.

### The GC B cell population in bladder-draining lymph nodes of UTI mice remains the same following MDR UPEC re-infection

Given the transient nature of the B cell response, we hypothesised that it could be boosted by administering a secondary inoculation of UPEC one week following the initial inoculation, due to increased antigen availability and uptake. At 4 weeks, C57BL/6 mice that had experienced a single UTI episode or were re-infected after one week with the same UPEC strain, had the same proportion of GC B cells in their bladder-draining lymph nodes (Figure 4). Both mouse groups had statistically increased GC B cell populations compared to naïve mice (*p* = 0.0002).

**Figure 4.**
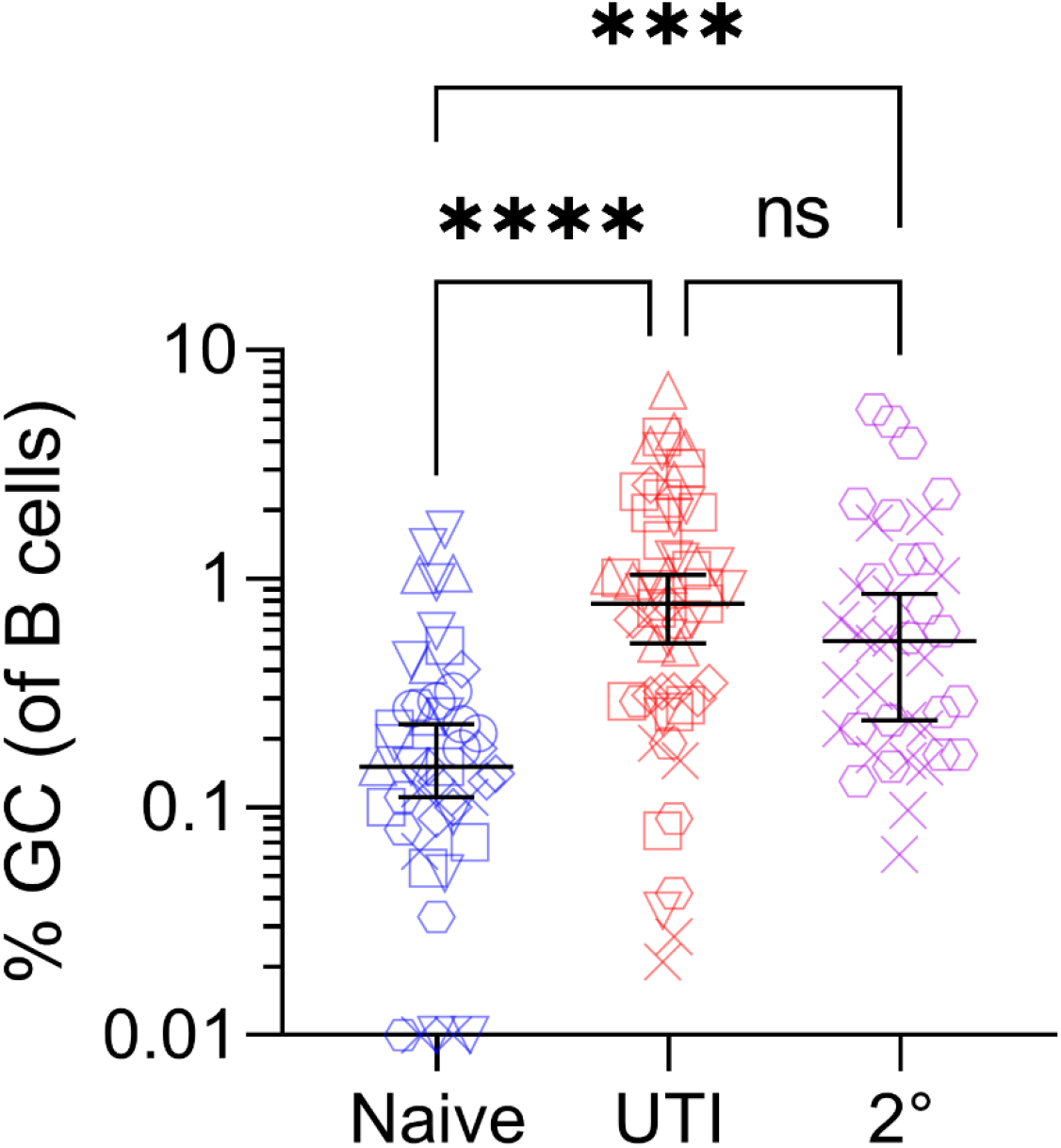
Secondary MDR UPEC inoculation after the first week of acute UTI does not increase the proportion of GC B cells present in local lymph nodes in C57BL/6 mice. Percentage of GC B cells of the total number of B cells detected at 4 weeks in bladder-draining lymph nodes of C57BL/6 naïve mice (*n* = 47), mice receiving a single dose of UTI inoculum (UTI: *n* = 57), or a secondary inoculation (2°) one week after the primary (*n* = 40). Group differences detected by Kruskal-Wallis test, bars represent median ±95% CI, **ns** not significant, **\*\*\*** *p <*0.001, **\*\*\*\****p <*0.0001).

## Discussion

In this study, we provide evidence consistent with the hypothesis that local humoral immune responses are generated during UTI in mice. Firstly, using *rag1*^*-/-*^ mice we conclude that T and/or B cells likely contribute to infection control, and importantly we show for the first time that GC B cell responses are generated in bladder draining lymph nodes. However, these responses appear to be relatively modest and transient in nature and were not boosted upon bladder reinfection. Given previous reports of bacteria-specific antibody generation during UTI in both mice and humans (25-27), our data suggests that at least some of the upstream T and B cell immune responses that lead to antibody production likely occur in local draining lymph nodes.

Transient GC B cell responses are not unusual in other infection settings, since they subside as the prevalence of antigen decreases (13). However, we observed that despite the return of GC B cell proportions to baseline by 7 wpi, detectable bacteria still remained in bladders. This implies that the local immune response is insufficient to eradicate bacteria. UPEC employ a variety of mechanisms to avoid clearance, including the formation of intracellular bacterial communities (IBC) and quiescent intracellular reservoirs (QIR) (28, 29), and blocking of TLR4 and cytokine signalling (17). Hence, we speculate that bacterial colonisation strategies in this model are able to survive, and perhaps even subvert, the host humoral immune response.

The magnitude of the GC B cell response varied substantially among individual mice. Reasons for this variability are unclear, but may have arisen from differences in innate immune response, which has previously been shown to affect infection outcomes (18, 30). High and transient levels of TNFα, IL-6, and IL-8 are associated with bacterial clearance, whereas sustained cytokine levels for prolonged periods of time are associated with tissue damage in the bladder (17, 18, 30). Pre-existing antibody titres against the infecting strain also affect the rate of bacterial clearance (31), suggesting that pre-existing antibodies to commensal organisms in “specific pathogen free” conditions might have influenced outcomes, although how this might induce variation between co-housed mice is unclear. Interestingly, secondary inoculation of mice did not lead to increased clearance rates compared to mice infected once, suggesting either that there was insufficient time for titres to develop or that the GC B cell response overall is not strong enough to produce meaningful antibody titres against the infecting UPEC strain EC958. The variation in responses could also be due to individual differences between mice, a common effect observed even in genetically identical mice (32).

The GC B cell response that we observed in bladder-draining lymph nodes is anecdotally weak compared to other models used in our groups (e.g. blood-stage *Plasmodium spp*. infection) (33), leading us to speculate that specific aspects of the bladder may contribute to this difference. For instance, the architecture of the bladder is structured in a way that is difficult for molecules to enter or leave (34, 35). During cystitis, the most important anatomical structures are superficial bladder cells that come into contact with UPEC and the contents of the bladder lumen and are held together with tight junctions. These cells are covered with plaques called uroplakins which add additional protection against toxic compounds in urine and pathogens (35, 36). Insufficient antigen uptake from the bladder to nearby lymph nodes would also affect antigen presentation and subsequent activation of adaptive immune cells. As increased bladder damage has been reported to lead to a stronger response (30, 37), we reasoned that administering a second UPEC inoculation, as performed in previous studies (15, 38, 39), might enhance antigen uptake and presentation. The modest GC B cell responses could theoretically also be the result of adaptive immune suppression via cytokines, which is a widely cited reason for a lack of a strong adaptive response in the bladder (8, 40-43). Early during the innate immune response to UPEC, IL-10 is secreted by mast cells to dampen inflammation and prevent bladder damage (40, 44). This in turn decreases DC activation, and CD4^+^ T cell and GC B cell responses in draining lymph nodes. Hence we speculate that one possible method for boosting antibody production during UTI might be to transiently block IL-10 signalling, as it has been shown that IL-10 knockout mice can more efficiently clear UPEC from their bladders in early infection and generate an increased UPEC-specific antibody titre (40).

Recently, advances have been made in understanding roles played by T cells during UTI (37, 38). A population of microbiota-dependent memory CD4^+^ T cells has been found to exist in bladder-draining lymph nodes, which respond to activation by DCs trafficking from the infected bladder (37). Interestingly, after secondary cystitis, there is preferential activation of Th2 cells within the bladder-draining lymph nodes by DCs, skewing the immune response to that of tissue repair (37). This process was found to be triggered by the exfoliation of bladder cells during infection, which upregulated a subset of these DCs (CD301b+ OX40L+). Skewing towards a Th2-dependent response was found to supress bacterial clearance, while amplifying the bladder’s tissue repair processes (37). Administration of a Th1-skewing adjuvant, however, was able to rectify this and again promote bacterial clearance (38). Taken together, these factors suggest that multiple host and pathogen factors contribute to the severity of infection and the type of immune response that occurs during UTI.

In conclusion, we demonstrated that a humoral response is generated locally during UTI that is important for infection control. The local GC B cell response is however relatively short-lived. Further research into the types of B cell subsets present and how the GC B cell response could be amplified by vaccination would be of benefit to the field. Boosting GC B cell responses with a range of adjuvants and/or immune-modulatory treatments could serve to improve protective immunity to UTI. Finally, shedding light on strategies for boosting adaptive immunity in UTI could pave the way to lowering the rate of recurrence for one of the most common bacterial infections in humans.

## Methods

### Bacterial strain and culture conditions

UPEC strain EC958 was used in this study as a reference multidrug resistant ST131 strain (45). To promote the expression of type 1 fimbriae (T1F), EC958 was routinely cultured from highly fimbriated stocks, which were incubated overnight statically at 37°C. Cultures were assessed for expression of T1F by yeast cell agglutination as previously described (46) and were fixed at the inoculation cell density and volume (47).

### Mouse UTI model and ethics

6–7-week-old female C57BL/6 and 8-17-week-old *rag1*^*-/-*^ mice were catheterised as previously described (48). Briefly, mice were anaesthetised by isoflurane inhalation and catheterised with ∼1-2 × 10^8^ CFU in 30 μL of UPEC. The prepared inoculum was deposited directly into the mouse bladder using a sterile catheter followed by immediate removal (48). A separate cohort of strain and aged-matched mice were used as controls and were not catheterised with bacteria. For innate immunity experiments, an additional cohort of mice were mock-catheterised with PBS. For experiments involving multiple inoculations, secondary inoculation mice were catheterised as described above and then after 1 week were catheterised again with the same dose of bacteria. C57BL/6 mice were sourced from the ARC (Animal Resource Centre, Western Australia) and *rag1*^*-/-*^ mice were kindly provided by Associate Professor Ashraful Haque, bred under QIMR-B ethics approval number A1503-601M. All UTI mouse work was conducted under QIMR-B ethics approval number A1703-600M and Queensland University of Technology OGTR approval number 1700000118. Mice were sacrificed at either 1 day, 4 weeks or 7 weeks post inoculation and the bladders, kidneys, and lymph nodes (lumbar aortic, medial iliac, mesenteric) were extracted.

### Flow cytometry and cell sorting

Bladders were incubated at 37°C for 1 hour in RPMI containing 1 mg/mL collagenase IV (Sigma, cat no. C1889-50MG, Darmstadt) and DNase I (Sigma, cat no. D4263-5VL, Darmstadt). Enzyme-treated bladders, and lymph nodes were mechanically disrupted with a 100 μm cell strainer (Falcon, cat no. 352360, Corning, NY) and resuspended in FACS buffer (1% FCS in PBS). 200 μL of each sample was plated onto 96-well plate (Falcon, cat no. 353077, Corning NY). Two draining lymph nodes were pooled per mouse. All samples were stained with Live/Dead Aqua (Invitrogen, L34965, Waltham, MA) and the respective antibody panels in Table 1. Samples were acquired on the Cytoflex S (Beckman-Coulter, Brea, CA) and BD Fortessa (BD Biosciences, Franklin Lakes, NJ), and data were processed using *FlowJo v10* (TreeStar, Franklin Lakes, NJ) software.

**Table 1.**
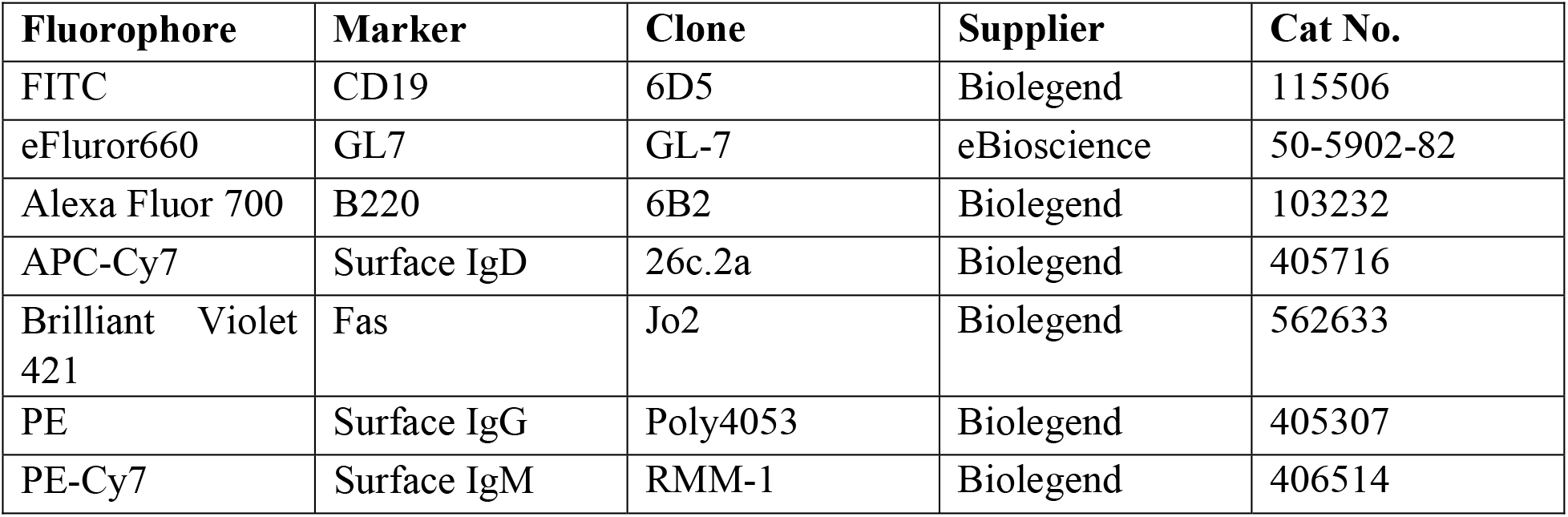
B cell antibody panel used in this study.

**Table 2.** Myeloid antibody panel used in this study.

| Fluorophore | Marker | Clone | Supplier | Cat No. |
| --- | --- | --- | --- | --- |
| FITC | Ly6C | HK1.4 | Biolegend | 128006 |
| PerCp-Cy5.5 | CD11b | M1/70 | Biolegend | 101228 |
| APC | CD11c | N418 | Biolegend | 117310 |
| Alexa Fluor 700 | B220 | RA3-6B2 | Biolegend | 103232 |
| Alexa Fluor 700 | TCR-β | H57-597 | Biolegend | 109224 |
| APC-Cy7 | Ly6G | 1A8 | Biolegend | 127624 |
| Pacific Blue | I/A-I/-E (MHC-II) | M5/114.15.2 | Biolegend | 107620 |
| PE | CD86 | GL-1 | Biolegend | 105008 |

### Quantification of bacterial viable cell numbers

Bacterial loads were quantified over time in urine, and at endpoint in infected mouse bladders. Urine samples were plated directly into a 96-well plate, bladders were homogenised in 50 μL of PBS using a Mini Beadbeater (BioSpec Products) before topping up to 1 mL with PBS and aliquoting 200 μL into the plate. Samples were serially diluted to 10^−4^ as previously described (48). 5 μL of each well for each sample was plated onto LB agar (1% tryptone, 0.5% yeast extract, 0.5% salt and 1.5% agar in Milli-Q water) in quadruplicate and then incubated overnight at 37°C. The following day, colonies were enumerated, and bacterial load was expressed as CFU/mL for urine or CFU/0.1g of tissue for bladders.

### Statistical Analysis

All statistical analysis was performed in *GraphPad Prism Version 9* software (GraphPad Software) and R(49). Unpaired non-parametric Mann-Whitney or Kruskal-Wallis tests were used to test for statistically significant differences between group medians of uninfected (naïve) and UTI groups. Statistical significance was set at *p* <0.05. Each unique symbol in Figures 1, 3, and 4 denotes an experimental repeat.

#### Statistical modelling of longitudinal urinalysis data

For analysis of longitudinal urine bacterial load and susceptibility to colonisation, a series of models were applied and tested in order to find the most appropriate model that fitted the nature of the data the best. Simple linear models were fitted with the log(CFU) as outcome and “Day “ and “Strain” as explanatory variables, where “Day” refers to the day post inoculation and “Strain” refers to mouse strain. The “lme” procedure from “nlme” package in R was then used to explore if a mixed model was needed, with “mouse” as a random effect, which would consider the potentially correlated nature of the data due to measurements taken on the same mouse over time. We also fitted a range of Generalised Additive Models (GAM) and Generalised Additive Mixed Models (GAMM). These models are similar to the above but instead of assuming a constant regression coefficient, they estimate a function(s) for the range of independent variables. Given the presence of a large number of zero values in the urine CFU data, zero inflated models were tested as well, as per Zuur et al. (50, 51). For these models both a Poisson and a Negative Binomial distribution were fitted, with the latter potentially accommodating a variance larger than the mean (i.e. overdispersion), at the expense of estimating one more parameter compared to the Poisson distribution. From this analysis, the model which fitted the data best (as judged by the Akaike’s information Criterion) was a Zero Inflated Negative Binomial Mixed model (ZINBMM). R Packages used in this analysis included: “nlme”, “lme4”, “mgcv”, “glmmTMB”, “pscl”, “lmtest” and “ggplot2” (52-58).

## Acknowledgements

This work was supported by a Faculty of Health Collaborative Seed grant from the Queensland University of Technology to M.T. and A.H. and in part, by a Clive and Vera Ramaciotti Health Investment grant (2017HIG0119) and a Georgina Sweet Award for Women in Quantitative Biomedical Science to M.T. S.H. is the recipient of an Australian Government Research Training Program (RTP) Scholarship. The authors would like to thank Jonathan Mauclair, Agustin Mercau and Taylah Anderson for technical assistance.

## Author Contributions

AH and MT designed the study and provided funding. SH conducted all experimental work and drafted the manuscript, figures, and tables. DV conducted statistical analyses of experimental data and contributed to drafting of the original manuscript. DV, AH, and MT edited the manuscript and figures. All authors contributed to the article and approved the submitted version.

## References

1. Foxman B. 2014. Urinary tract infection syndromes: occurrence, recurrence, bacteriology, risk factors, and disease burden. Infect Dis Clin North Am 28:1–13.

2. Foxman B, Brown P. 2003. Epidemiology of urinary tract infections: transmission and risk factors, incidence, and costs. Infect Dis Clin North Am 17:227–41.

3. Totsika M, Beatson SA, Sarkar S, Phan MD, Petty NK, Bachmann N, Szubert M, Sidjabat HE, Paterson DL, Upton M, Schembri MA. 2011. Insights into a multidrug resistant Escherichia coli pathogen of the globally disseminated ST131 lineage: genome analysis and virulence mechanisms. PLoS One 6:e26578.

4. Geerlings SE. 2016. Clinical Presentations and Epidemiology of Urinary Tract Infections. Microbiol Spectr 4.

5. Engel D, Dobrindt U, Tittel A, Peters P, Maurer J, Gutgemann I, Kaissling B, Kuziel W, Jung S, Kurts C. 2006. Tumor necrosis factor alpha- and inducible nitric oxide synthase-producing dendritic cells are rapidly recruited to the bladder in urinary tract infection but are dispensable for bacterial clearance. Infect Immun 74:6100–7.

6. Mora-Bau G, Platt AM, van Rooijen N, Randolph GJ, Albert ML, Ingersoll MA. 2015. Macrophages Subvert Adaptive Immunity to Urinary Tract Infection. PLoS Pathog 11:e1005044.

7. Scherberich JE, Hartinger A. 2008. Impact of Toll-like receptor signalling on urinary tract infection. Int J Antimicrob Agents 31 Suppl 1:S9–14.

8. Choi HW, Abraham SN. 2016. Why Serological Responses during Cystitis are Limited. Pathogens 5.

9. Lacerda Mariano L, Ingersoll MA. 2020. The immune response to infection in the bladder. Nat Rev Urol 17:439–458.

10. Chamoun MN, Sullivan MJ, Goh KGK, Acharya D, Ipe DS, Katupitiya L, Gosling D, Peters KM, Sweet MJ, Sester DP, Schembri MA, Ulett GC. 2020. Restriction of chronic Escherichia coli urinary tract infection depends upon T cell-derived interleukin-17, a deficiency of which predisposes to flagella-driven bacterial persistence. FASEB J 34:14572–14587.

11. Jones-Carson J, Balish E, Uehling DT. 1999. Susceptibility of immunodeficient gene-knockout mice to urinary tract infection. J Urol 161:338–41.

12. Thumbikat P, Waltenbaugh C, Schaeffer AJ, Klumpp DJ. 2006. Antigen-specific responses accelerate bacterial clearance in the bladder. J Immunol 176:3080–6.

13. Mesin L, Ersching J, Victora GD. 2016. Germinal Center B Cell Dynamics. Immunity 45:471–482.

14. Kantele A, Mottonen T, Ala-Kaila K, Arvilommi HS. 2003. P fimbria-specific B cell responses in patients with urinary tract infection. J Infect Dis 188:1885–91.

15. Hopkins WJ, Gendron-Fitzpatrick A, Balish E, Uehling DT. 1998. Time course and host responses to Escherichia coli urinary tract infection in genetically distinct mouse strains. Infect Immun 66:2798–802.

16. Hopkins WJ, Uehling DT, Balish E. 1987. Local and systemic antibody responses accompany spontaneous resolution of experimental cystitis in cynomolgus monkeys. Infect Immun 55:1951–6.

17. Billips BK, Forrestal SG, Rycyk MT, Johnson JR, Klumpp DJ, Schaeffer AJ. 2007. Modulation of host innate immune response in the bladder by uropathogenic Escherichia coli. Infect Immun 75:5353–60.

18. Hannan TJ, Mysorekar IU, Hung CS, Isaacson-Schmid ML, Hultgren SJ. 2010. Early severe inflammatory responses to uropathogenic E. coli predispose to chronic and recurrent urinary tract infection. PLoS Pathog 6:e1001042.

19. Schaale K, Peters KM, Murthy AM, Fritzsche AK, Phan MD, Totsika M, Robertson AA, Nichols KB, Cooper MA, Stacey KJ, Ulett GC, Schroder K, Schembri MA, Sweet MJ. 2016. Strain- and host species-specific inflammasome activation, IL-1beta release, and cell death in macrophages infected with uropathogenic Escherichia coli. Mucosal Immunol 9:124–36.

20. Schilling JD, Mulvey MA, Vincent CD, Lorenz RG, Hultgren SJ. 2001. Bacterial invasion augments epithelial cytokine responses to Escherichia coli through a lipopolysaccharide-dependent mechanism. J Immunol 166:1148–55.

21. Schwartz DJ, Chen SL, Hultgren SJ, Seed PC. 2011. Population dynamics and niche distribution of uropathogenic Escherichia coli during acute and chronic urinary tract infection. Infect Immun 79:4250–9.

22. Sivick KE, Schaller MA, Smith SN, Mobley HL. 2010. The innate immune response to uropathogenic Escherichia coli involves IL-17A in a murine model of urinary tract infection. J Immunol 184:2065–75.

23. Ulett GC, Totsika M, Schaale K, Carey AJ, Sweet MJ, Schembri MA. 2013. Uropathogenic Escherichia coli virulence and innate immune responses during urinary tract infection. Curr Opin Microbiol 16:100–7.

24. Pieper K, Grimbacher B, Eibel H. 2013. B-cell biology and development. J Allergy Clin Immunol 131:959–71.

25. Sarkissian CA, Alteri CJ, Mobley HLT. 2019. UTI patients have pre-existing antigen-specific antibody titers against UTI vaccine antigens. Vaccine 37:4937–4946.

26. Szijarto V, Guachalla LM, Visram ZC, Hartl K, Varga C, Mirkina I, Zmajkovic J, Badarau A, Zauner G, Pleban C, Magyarics Z, Nagy E, Nagy G. 2015. Bactericidal monoclonal antibodies specific to the lipopolysaccharide O antigen from multidrug-resistant Escherichia coli clone ST131-O25b:H4 elicit protection in mice. Antimicrob Agents Chemother 59:3109–16.

27. Thomas V, Shelokov A, Forland M. 1974. Antibody-coated bacteria in the urine and the site of urinary-tract infection. N Engl J Med 290:588–90.

28. Justice SS, Hung C, Theriot JA, Fletcher DA, Anderson GG, Footer MJ, Hultgren SJ. 2004. Differentiation and developmental pathways of uropathogenic Escherichia coli in urinary tract pathogenesis. Proc Natl Acad Sci U S A 101:1333–8.

29. Mysorekar IU, Hultgren SJ. 2006. Mechanisms of uropathogenic Escherichia coli persistence and eradication from the urinary tract. Proc Natl Acad Sci U S A 103:14170–5.

30. Yu L, O’Brien VP, Livny J, Dorsey D, Bandyopadhyay N, Colonna M, Caparon MG, Roberson ED, Hultgren SJ, Hannan TJ. 2019. Mucosal infection rewires TNFa signaling dynamics to skew susceptibility to recurrence. Elife 8:e46677.

31. Hopkins WJ, Uehling DT. 1995. Resolution time of Escherichia coli cystitis is correlated with levels of preinfection antibody to the infecting Escherichia coli strain. Urology 45:42–6.

32. Voelkl B, Altman NS, Forsman A, Forstmeier W, Gurevitch J, Jaric I, Karp NA, Kas MJ, Schielzeth H, Van de Casteele T, Würbel H. 2020. Reproducibility of animal research in light of biological variation. Nat Rev Neurosci 21:384–393.

33. James KR, Soon MSF, Sebina I, Fernandez-Ruiz D, Davey G, Liligeto UN, Nair AS, Fogg LG, Edwards CL, Best SE, Lansink LIM, Schroder K, Wilson JAC, Austin R, Suhrbier A, Lane SW, Hill GR, Engwerda CR, Heath WR, Haque A. 2018. IFN Regulatory Factor 3 Balances Th1 and T Follicular Helper Immunity during Nonlethal Blood-Stage <em>Plasmodium</em> Infection. The Journal of Immunology 200:1443–1456.

34. Lewis SA. 2000. Everything you wanted to know about the bladder epithelium but were afraid to ask. Am J Physiol Renal Physiol 278:F867–74.

35. Wu J, Miao Y, Abraham SN. 2017. The multiple antibacterial activities of the bladder epithelium. Ann Transl Med 5:35.

36. Hill WG. 2015. Control of urinary drainage and voiding. Clin J Am Soc Nephrol 10:480–92.

37. Wu J, Hayes BW, Phoenix C, Macias GS, Miao Y, Choi HW, Hughes FM, Jr., Todd Purves J, Lee Reinhardt R, Abraham SN. 2020. A highly polarized TH2 bladder response to infection promotes epithelial repair at the expense of preventing new infections. Nat Immunol 21:671–683.

38. Wu J, Bao C, Reinhardt RL, Abraham SN. 2021. Local induction of bladder Th1 responses to combat urinary tract infections. Proc Natl Acad Sci U S A 118.

39. Hjelm EM. 1984. Local cellular immune response in ascending urinary tract infection: occurrence of T-cells, immunoglobulin-producing cells, and Ia-expressing cells in rat urinary tract tissue. Infect Immun 44:627–32.

40. Chan CY, St John AL, Abraham SN. 2013. Mast cell interleukin-10 drives localized tolerance in chronic bladder infection. Immunity 38:349–59.

41. Chan CY, St John AL, Abraham SN. 2012. Plasticity in mast cell responses during bacterial infections. Current Opinion in Microbiology 15:78–84.

42. Abraham SN, Miao Y. 2015. The nature of immune responses to urinary tract infections. Nature Reviews Immunology 15:655–63.

43. Mora-Bau G, Platt AM, van Rooijen N, Randolph GJ, Albert ML, Ingersoll MA. 2015. Macrophages Subvert Adaptive Immunity to Urinary Tract Infection. PLoS Pathogens 11:e1005044.

44. Acharya D, Sullivan MJ, Duell BL, Goh KGK, Katupitiya L, Gosling D, Chamoun MN, Kakkanat A, Chattopadhyay D, Crowley M, Crossman DK, Schembri MA, Ulett GC. 2019. Rapid Bladder Interleukin-10 Synthesis in Response to Uropathogenic Escherichia coli Is Part of a Defense Strategy Triggered by the Major Bacterial Flagellar Filament FliC and Contingent on TLR5. mSphere 4.

45. Forde BM, Ben Zakour NL, Stanton-Cook M, Phan MD, Totsika M, Peters KM, Chan KG, Schembri MA, Upton M, Beatson SA. 2014. The complete genome sequence of Escherichia coli EC958: a high quality reference sequence for the globally disseminated multidrug resistant E. coli O25b:H4-ST131 clone. PLoS One 9:e104400.

46. Totsika M, Kostakioti M, Hannan TJ, Upton M, Beatson SA, Janetka JW, Hultgren SJ, Schembri MA. 2013. A FimH inhibitor prevents acute bladder infection and treats chronic cystitis caused by multidrug-resistant uropathogenic Escherichia coli ST131. Journal of Infectious Diseases 208:921–8.

47. Hung CS, Dodson KW, Hultgren SJ. 2009. A murine model of urinary tract infection. Nature Protocols 4:1230–43.

48. Hannan TJ, Hunstad DA. 2016. A Murine Model for Escherichia coli Urinary Tract Infection. Methods Mol Biol 1333:159–75.

49. R Core Team. 2022. R: A language and environment for statistical computing., R Foundation for Statistical Computing, Vienna, Austria.

50. Zuur AF, Ieno EN. 2016. Beginner’s guide to zero-inflated models with R.

51. Zuur AF, Savaliev AA, Ieno EN. 2012. Zero inflated models and generalized linear mixed models with R. Highland Statistics, Newburgh, Scotland.

52. Pinheiro J BD, R Core Team. 2022. nlme: Linear and Nonlinear Mixed Effects Models., vR package version 3.1–158.

53. Bates D, Mächler M, Bolker B, Walker S. 2015. Fitting Linear Mixed-Effects Models Usinglme4. Journal of Statistical Software 67:1–48.

54. Wood SN. 2011. Fast stable restricted maximum likelihood and marginal likelihood estimation of semiparametric generalized linear models. Journal of the Royal Statistical Society Series B-Statistical Methodology 73:3–36.

55. Brooks M, Kristensen K, van Benthem K, Magnusson A, Berg CW, Nielsen A, Skaug H, Mächler M, Bolker B. 2017. glmmTMB Balances Speed and Flexibility Among Packages for Zero-inflated Generalized Linear Mixed Modeling. R Journal 9:378–400.

56. Zeileis A, Kleiber C, Jackman S. 2008. Regression Models for Count Data in R. Journal of Statistical Software 27:1 –25.

57. Zeileis A. HT. 2002. Diagnostic Checking in Regression Relationships. https://CRAN.R-project.org/doc/Rnews/. Accessed 15/02.

58. Wickham H. 2016. ggplot2: Elegant Graphics for Data Analysis, Springer-Verlag New York.

